# Evidence for a push-pull interaction between superior colliculi in monocular dynamic vision mode

**DOI:** 10.1101/2024.05.06.592678

**Authors:** Rita Gil, Mafalda Valente, Francisca F. Fernandes, Noam Shemesh

## Abstract

Visual perception can operate in two distinct vision modes - static and dynamic - that have been associated with different neural activity regimes in the superior colliculus (SC). The static vision mode (low flashing frequencies) is associated with strong SC activation modulated by cortical gain and inhibitory intertectal effects, while the dynamic vision mode (high flashing frequencies) evokes the continuity illusion, with associated suppression of SC neural activity. However, the pathway-wide mechanisms underpinning the dynamic vision mode remain poorly understood, especially in terms of corticotectal and tectotectal feedback. Here, we harness rat functional MRI combined with brain lesions to investigate whole-pathway interactions in the dynamic vision mode. In the SC, we find contralateral suppression of activity opposing positive ipsilateral neural activation upon monocular visual stimulation in the dynamic vision mode. A cortical amplification effect was confirmed for both static and dynamic vision modes through cortical lesions, while further lesioning ipsilateral SC led to a boost in the contralateral negative signals, suggesting an active push-pull interaction between ipsilateral and contralateral SCs during the dynamic vision mode regime. This push-pull interaction is specific to the dynamic vision mode; in the static vision mode, both SCs show similar response polarities. These results highlight hitherto unreported frequency-dependent modulations in the tectotectal pathway and further challenge the contemporary notion that intertectal connections solely serve as reciprocal inhibitory mechanisms for avoiding visual blur during saccade occurrence.

**One Sentence Summary:** Opposing signals between superior colliculi in the dynamic vision mode suggest an active push-pull interaction within the tectotectal commissural pathway.

## Introduction

The rapid assessment of the surrounding environment and appropriate behavioural response to external stimuli (e.g. to approach, avoid or ignore) is key for animal survival^1^. The continuity illusion is a perceptual feature enabling the brain to interpret moving surroundings smoothly and detect new emerging features in a continuously stable environment. When the frequency of the light reaching the retina is low, each flash is perceived as unique and the system operates in static vision mode; if the stimuli is with a temporal frequency above the flicker fusion frequency (FFF), the system operates in the dynamic vision mode where the stimuli are fused to an illusory continuous percept.

The superior colliculus (SC) is an important relay for sensory pathways (e.g. visual, auditory and somatosensory) with several outputs towards motor areas^2^ and it is known to play critical roles in multimodal integration and rapid sensorimotor transformation^3–5^. Cells in the SC superficial visual layers (sSC) are imperative for visual saliency detection^3,6–8^, i.e. stimulus appearance/disappearance or movement (important for innate defensive responses such as escape or camouflage from predators). Importantly, the SC activity was found to strongly depend on flashing stimulus frequency associated with the encoding of the response habituation (RH) phenomenon^9,10^ characterised by response decrements, both in total number of spikes^6^ and local field potentials (LFPs) N1-P1 amplitude attenuation^11^, to repeated stimuli presentations. RH has been thought of as a form of short-term memory for familiar versus novel information based on a dynamic adjustment of response thresholds. This phenomenon linked SC also with novelty detection, and was shown to be underpinned by multiple inhibitory feedback mechanisms^12,13^ occurring between and within the two SCs: the coactivation of excitatory and inhibitory neurons leads to a long-lasting inhibition that would block responses to subsequent stimulus at sufficiently high frequencies. In higher-order primates and humans, the higher visual cortical areas have been more implicated with visual perception. However, the role of SC as “novelty detector” along with other rodent collicular properties usually associated with cortical regions of primates (such as orientation selectivity^14^), suggest a more prominent role of the SC alongside the visual cortex (VC) in rodent visual processing. A recent study encompassing behavioural readouts, functional MRI (fMRI), and electrophysiology in the rat animal model found the most marked correlation with flashing stimuli frequency occurring at the level of the SC, where transitions between activation and suppression states matched the behaviourally reported flicker fusion frequency (FFF) threshold, where the shifts from static to dynamic vision modes takes place^15^. Interestingly, the SC fMRI signals also revealed the expected characteristics of “novelty detection”, i.e. strong flash-induced positive signals in low frequency stimulation (representing the novelty introduced by each flash) and onset/offset positive peaks surrounding the stimulation period in the dynamic vision mode (representing the novelty of brightness changes since the stimulus is perceived as a continuous light).

The SC is positioned strategically in the midbrain, and its high connectivity^2–5^ with other areas suggest a potential role for inter-area interactions that could contribute to the encoding and operation of the vision modes. In the rodent brain, around 85-90% of retinal ganglion cells (RGCs) project directly to the sSC. As retinal evoked potentials can still track individual light flashes during the dynamic vision mode^16,17^, the SC activity modulation is impacted by feedback arriving to the region. Several feedback connections are known, mainly from the ipsilateral primary visual cortex (V1) through corticotectal connections^3,15,18–20^ and from the contralateral SC (cSC) through intertectal connections^4,5,21^. Corticotectal connections project to all layers of the SC and are known to be aligned with retinal inputs. These connections on tectal cells are thought to exert a gain control effect: sSC neurons inherit feature selectivity from retinal hardwired circuits and modulate their response magnitude depending on cortical input that may be subject to modulation by, for example, the animal’s internal state, attention and learning^3,15,18–20^. Reduced responsiveness in many sSC neurons has been reported after inactivation of corticotectal feedback connections, either by cortical removal, pharmacological depression, optogenetic silencing or cool down^19,20,22–25^. Tectotectal connections between the two superior colliculi are also known to play important roles in visual-orienting behaviours^26^: the two SCs are connected through commissural fibres thought to be responsible for a reciprocal intertectal inhibitory effect^18,27–31^ related to surround inhibition and to suppression of visual inputs during saccades which would otherwise lead to visual blur^1^. The existence of direct inhibitory connections between the cSC and ipsilateral SC (iSC) was confirmed by intracellular recordings of inhibitory postsynaptic potentials from collicular cells in response to electrical stimulation of the cSC^30^. Although most electrophysiological studies support a predominantly inhibitory intertectal effect, evoked activation in several tectal cells after stimulation of intertectal fibres has also been reported^7,26,27^.

Studies of corticotectal and tectotectal connections were performed in the static vision mode, where SC activation is expected to occur following a visual stimulus. However, how the two colliculi interact between themselves and with other areas during the dynamic vision mode is still poorly understood. Here, we harness fMRI, which allows for a whole-pathway perspective, and brain lesions in key visual pathway structures, to investigate collicular interactions and the impact of corticotectal and tectotectal feedback in the observed SC neural suppression in the dynamic vision mode. A monocular stimulation regime was used to actively isolate activity derived from the visual stimulus into one hemisphere and to allow the investigation of interhemispheric tectal connections. We discover a push-pull interaction between the two SCs during the dynamic vision mode upon monocular stimulation, pointing at the role of tectotectal feedback beyond general reciprocal inhibitory tectotectal feedback.

## Methods

All animal care and experimental procedures were carried out according to the European Directive 2010/63 and pre-approved by the competent authorities, namely, the Champalimaud Animal Welfare Body and the Portuguese Direcção-Geral de Alimentação e Veterinária (DGAV).

In this study, n = 27 adult Long-Evans rats (n = 19 females) were used. The animals had *ad libitum* access to food and water and were kept under normal 12h/12h light/dark cycle. Rats were randomly split into three groups (**Figure 1A**): The fMRI healthy group consisted of 7 rats (n = 4 females): 14.9 ± 3.9 weeks old, weighing 349.1 ± 87.5 g on average. The fMRI V1 lesioned group consisted of 8 rats (n = 3 females): 16.8 ± 0.9 weeks old, weighing 357.5 ± 57.9 g on average. The fMRI V1+iSC lesioned group consisted of 7 rats (n = 6 females): 24.7 ± 7.7 weeks old, weighing 306.3 ± 31.6 g on average. A fourth group where the iSC was lesioned consisted of 5 rats (n = 5 females): 22.8 ± 3.2 weeks old, weighing 400.6 ± 24.1 g on average.

**Figure 1:**
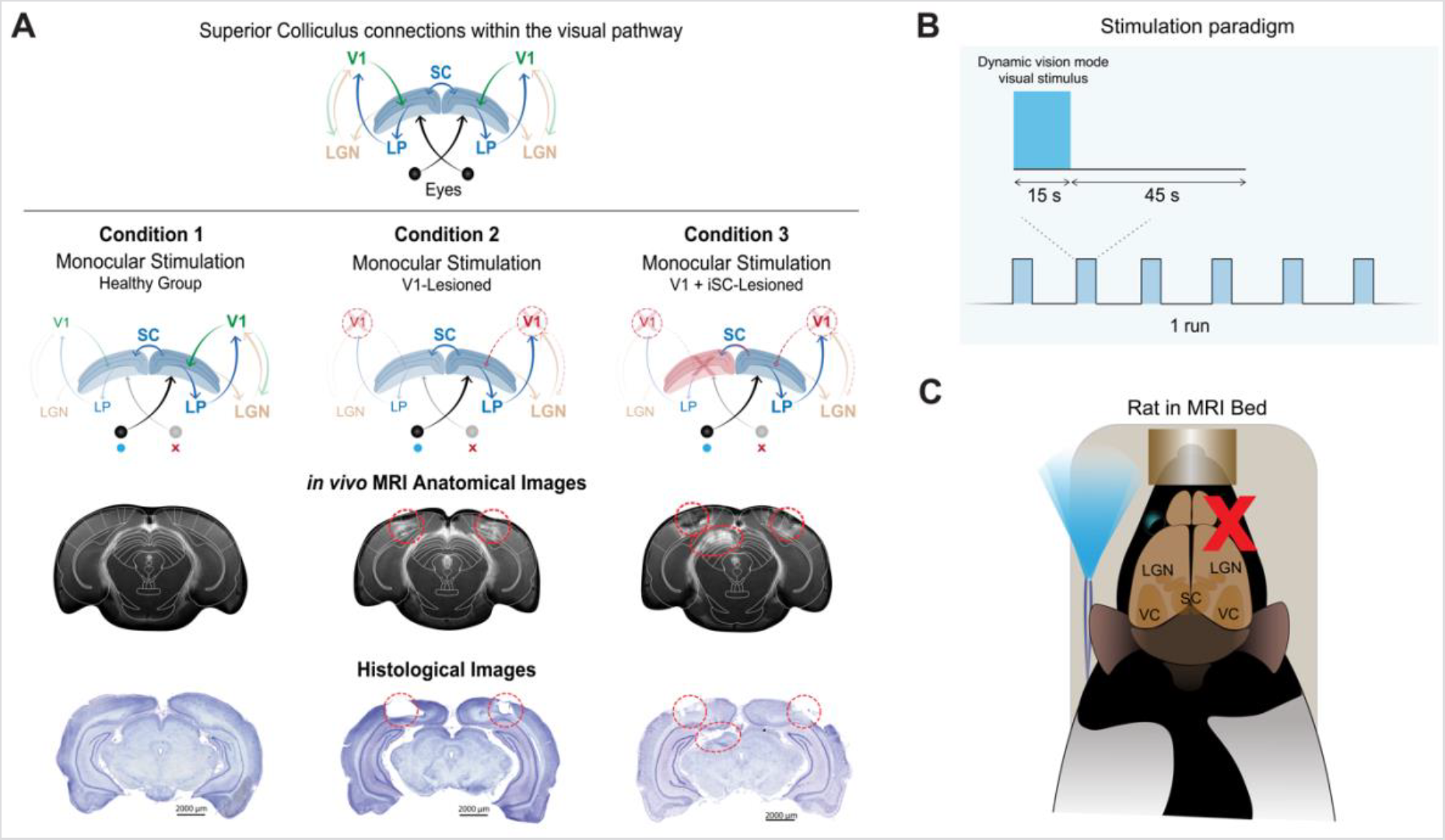
Experiment schematics. **(A)** Different tested conditions: Healthy group where no lesions were performed; V1-lesioned group where corticotectal feedback is reduced due to bilateral lesions in the V1; and V1+iSC-lesioned group where corticotectal and tectotectal feedback is reduced through bilateral lesions in the V1 combined with unilateral lesions in the iSC. MRI and histological images confirm the correct placement of the lesions for specific feedback reduction; **(B)** Stimulation paradigm consisting of 15 s of a high frequency flashing stimulus (frequency = 25 Hz) to induce the dynamic vision mode, followed by 45 s of rest. Each experimental run consisted of six repetitions of the described stimulation block; **(C)** Schematic of the animal in the MR bed with the LED tip placed in front of one eye while the other eye was covered for efficient monocular stimulation. V1 - primary visual cortex; iSC - ipsilateral superior colliculus.

### Ibotenic Acid Lesions and Perfusion

Animals were injected with an ibotenic acid (Abcam, Romania) solution (excitotoxic agent, 1mg/100 µL) either bilaterally in the V1 (n = 8), unilaterally in the iSC (n = 5), or simultaneously bilaterally in the V1 and the iSC (n = 7), to investigate the effect of cortical or tectal feedback projections in the cSC activity. Lesions performed with ibotenic acid are expected to predominantly target cholinergic neurons, while leaving the vasculature and passing fibres on that region intact^32,33^. The acid was injected using a Nanojet II (Drummond Scientific Company). Animals were anaesthetised with isoflurane (anaesthesia induced at 5% concentration and maintenance below 3% in medical air), and a scalpel incision was made along the midline of the skull, the skin retracted and the soft tissue cleaned from the skull with blunt forceps.

Coordinates for the craniotomies and number of injections required were determined for each individual animal based on T_2_-weighted anatomical images acquired before the surgery. Injections were made in a maximum of 5 different Anterior-Posterior (AP) coordinates, with 1 injection site for the first AP coordinate (2 pulses of injection) and 2 for the following (4 pulses of injection each), for full coverage of the V1. For full coverage of the SC, 3 injections were performed in different AP coordinates, each with 3 medio-lateral (ML) injection sites (vertical insertion, 2 pulses of injection each). For the second and third AP coordinate an extra injection was performed at a 20º angle with the vertical (2 pulses of injection). Each injection pulse was administered at a rate of 23 nL/s, waiting 2-3 s between pulses and each pulse consisted of 32 nL for the V1 and 23 nL for the SC injections. The waiting time before removing the injection pipette after the last pulse was 10 min. The craniotomies were then covered with Kwik-Cast™ (World Precision Instruments, USA) and the scalp was sutured. Before the animals recovered from surgery, they were injected, subcutaneously, with 5 mg/Kg body weight of carprofen (Rimadyl ®, Zoetis, U.S.A). Following surgery, the animals were allowed to recover for 7-9 days to avoid MRI acquisition artefacts due to potential inflammation and mechanical damage from the pipette.

For post-mortem confirmation of the lesions (after MRI scanning), the animals were overdosed with pentobarbital (100 mg/kg of body weight, intraperitoneal injection) and perfused transcardially with a solution of 4% PFA for a period of 12-24 h before slicing for further microscopy imaging (**Figure 1A**).

### Visual Set-Up and Paradigm for fMRI Acquisitions

The flashing stimulation was performed at a high frequency (25 Hz) known to induce the dynamic vision mode^15^, and the flash duration was set to 10 ms. The stimulation paradigm consisted of a 15 s stimulation period interleaved with 45 s rest periods (**Figure 1B**) and this cycle was repeated 6 times in each MRI acquisition run.

The stimulated eye of the animal was hydrated at the start of acquisition with ophthalmic gel (Vidisic gel Bausch + Lomb, Portugal) and the tip of an optic fibre connected to a blue LED (λ = 470 nm and I = 8.1×10^-1^ W/m^2^) were placed horizontally in front of it, spaced up to 1 cm from the animal (**Figure 1C**). To achieve a successful monocular stimulation regime, the other eye was covered with an opaque gel (Bepanthen augen und nasensalbe) and a piece of black polyurethane-coated nylon fabric to avoid any light from entering. The blue LED was connected to an Arduino MEGA2560 receiving triggers from the MRI scanner and was used to generate square pulses of light. Given the position of the optic fibre relative to the stimulated eye and the fact that the blue light reflected inside the MRI bore, the entire field of view of the animal was considered to be covered.

### Animal Preparation for fMRI Acquisition

Anesthesia was induced with 5% isoflurane (Vetflurane, Virbac, France) in medical air for 2 min, and the animals were then weighed and transferred to the MRI bed. Sedation began through an injection of a subcutaneous bolus (0.05 mg/kg) of medetomidine solution (1:10 dilution in saline of 1 mg/ml medetomidine solution - Vetpharma Animal Health, S.L., Spain), 5 min after the initial isoflurane induction. Isoflurane was gradually reduced to 0% during the following 10 min, whereupon a constant infusion of medetomidine (0.1 mg/kg/h) was initiated via a syringe pump (GenieTouch, Kent Scientific, Torrington, Connecticut, USA). For the rest of the MRI session, animals remained only under continuous medetomidine sedation. To ensure sufficient isoflurane washout, the fMRI acquisitions were started 30 min after medetomidine bolus injection. In the end of each MRI session, the medetomidine sedation was reverted by injecting an equal volume of the initial bolus of a 5 mg/ml solution of atipamezole hydrochloride (Vetpharma Animal Health, S.L., Spain) diluted 1:10 in saline. During the fMRI experiments, animals breathed a mixture of 95% oxygen and 5% medical air.

The animal’s temperature was constantly monitored with a rectal temperature optic fibre probe (SA Instruments, Inc., Stony Brook, New York, USA) and was kept at 36.5 ± 1.0 ºC via water circulating through a heat pad placed underneath the animal. Respiratory rate was measured using a pillow sensor (SA Instruments Inc., Stony Brook, USA) and was kept at 89.1 ± 17.7 breaths/min for the entire fMRI session.

### MRI Acquisitions

The experiments were conducted on a 9.4T Bruker Biospec MRI scanner (Bruker, Karlsruhe, Germany) equipped with an AVANCE III HD console, a gradient unit capable of producing pulsed field gradients of up to 660 mT/m isotropically with a 120 μs rise time, and ParaVision®6.0.1 software. An 86 mm quadrature coil was used for radiofrequency transmission and a 4-element array cryoprobe^34^ (Bruker, Fallanden, Switzerland) was used for signal reception.

For correct slice placement along the visual pathway, an anatomical T_2_-weighted Rapid Acquisition with Refocused Echoes (RARE) sequence was first acquired (TR/TE = 1600 / 36 ms, RARE factor = 8, Echo spacing = 9 ms; Averages = 3; FOV = 18 × 16 mm_2_, in-plane resolution = 168 × 150 μm^2^, slice thickness = 800 μm, t_acq_= 1 min 3 s). In a subset of animals, the anatomical acquisition was acquired with slightly different parameters: TR/TE = 3000 / 40 ms, RARE factor = 12, Echo spacing = 6 ms; Averages = 2; FOV = 18 × 16. mm^2^, in-plane resolution = 168 × 150 μm^2^, slice thickness = 800 μm, t_acq_= 48 s.

The functional MRI were acquired using a Spin-Echo Echo-Planar Imaging (SE-EPI) sequence (TR/TE= 1500 / 40 ms, Partial Fourier Factor (pFT) = 1.5, FOV = 18 x 16.1 mm^2^, in-plane resolution = 269 x 268 μm^2^, slice thickness = 1.5 mm, 8 slices, t_acq_= 6 min 50 s).

### fMRI data Analysis

The data was analysed using MATLAB (The Mathworks, Natick, MA, USA, v2016a and v2018b). A general linear model (GLM) analysis was conducted voxelwise along with a region of interest (ROI) analysis based on anatomy to gain insight into temporal dynamics of activation profiles.

#### GLM Analysis

Pre-processing steps included manual outlier removal through a spline interpolation taking the entire time course (<0.0005% of the data were identified as outliers), slice-timing correction (using a sinc-interpolation) followed by head motion correction (using a mutual information algorithm). Data was then co-registered to the T_2_-weighted anatomical images, normalised to a reference animal and smoothed using a 3D Gaussian isotropic kernel with full width half-maximum of 0.268 mm. A double gamma HRF peaking at 0.2 s was convolved with the stimulation paradigm to obtain an experimental regressor of the design matrix peaking at 1.75 s (around the time fMRI temporal profiles peaked).

From the dynamic vision mode contralateral time profiles, it became apparent that the cSC responses were multi-periodic (c.f. Results) and could be decomposed into three distinct periods: onset, ON and offset. Therefore, the three periods were used in a second GLM as 3 independent regressors after being convolved with a similar HRF peaking at 0.2 s. The onset regressor was designed to peak 1.75 s after the stimulation started; the ON regressor peaked 6.25 s after stimulation started; and the offset regressor peaked 3.25 s after the stimulation finished.

A fixed-effects group analysis was run independently for each group and for the difference between two groups. Resulting t-value maps were thresholded with p < 0.005 and a minimum cluster size of 10 voxels, and were cluster-FDR corrected at p < 0.005, both when analysing each group and the difference between groups.

#### ROI Analysis

The 6^th^ Edition of Paxinos & Franklin’s rat brain atlas^35^ was used for manual ROI delineation. The individual normalised fMRI runs were detrended with a 2^nd^ degree polynomial fit to the first resting period and the final 10 s of the subsequent resting periods to remove low frequency trends. The detrended data were then converted into percent signal change relative to baseline. For each run, the six individual cycles were separated and the averaged response was calculated within each ROI (along with the standard error of the mean), first at the animal level and afterward at the group level. Average signals per animal were used for statistical analysis.

#### Temporal signal-to-noise ratio (tSNR) Calculation

The tSNR was calculated to investigate the effect of the lesions in the ROI quantifications (**Figure S1A-B**). The ROIs drawn for the cSC and contralateral LGN during the ROI analysis step were used to calculate tSRN values. For each animal, the voxelwise mean and signal standard deviation during rest periods was calculated. The tSNR values were then obtained by dividing the mean signal by the standard deviation and the average values for each animal were calculated. Finally, for each group, the animal tSNRs for each ROI were averaged and the standard error of the mean was calculated.

## Results

### Opposing tectal responses in the monocular dynamic vision mode

We first examined the overall temporal profile of fMRI signals in ROIs placed along the visual pathway. Monocular stimulation at 25 Hz evoked strong activity in both cortical and subcortical regions of the visual pathway (**Figures 2 and S1**). Raw entire time profiles for the different ROIs do not exhibit habituation along the different stimulation blocks and further confirm the high data quality (**Figure S1C**). At the subcortical SC level, cSC responses could be clearly be divided into three distinct periods: two positive “peaks”, at the beginning and after the end of the visual stimulation period (hereafter referred to as onset and offset peaks, respectively, dashed arrows in **Figure 2A**), and a strong negative fMRI signal in-between (hereafter referred to as the ON period, solid arrow in **Figure 2A**). Unlike the strong negative signal during the ON period in the cSC, the iSC evidenced positive fMRI responses with less clear (if any) distinction between onset/ON/offset periods. Still at the subcortical level, positive signals were noted in lateral geniculate thalamic nucleus (LGN) for both hemispheres (with reduced percent signal change for the ipsilateral hemisphere, as expected from the monocular nature of the visual stimulus). Although not as prominent as in SC, the three periods could be somewhat discerned in the contralateral LGN responses (**Figure 2A**, middle panel, black arrows). At the cortical level, monocular stimulation induced strong bilateral negative signals in the V1, with stronger negative responses observed in the contralateral hemisphere. The three periods observed in the contralateral responses at the subcortical level were not distinguishable in these cortical responses. Although an onset was not evident, the peak cortical signal coincided with the subcortical ON periods (filled black arrow in the V1 time profile of **Figure 2A** at ∼6 s after stimulus started), and a more conventional post-stimulation signal (in this case, overshoot) was observed (dashed black arrow in the V1 time profile of **Figure 2A**).

**Figure 2:**
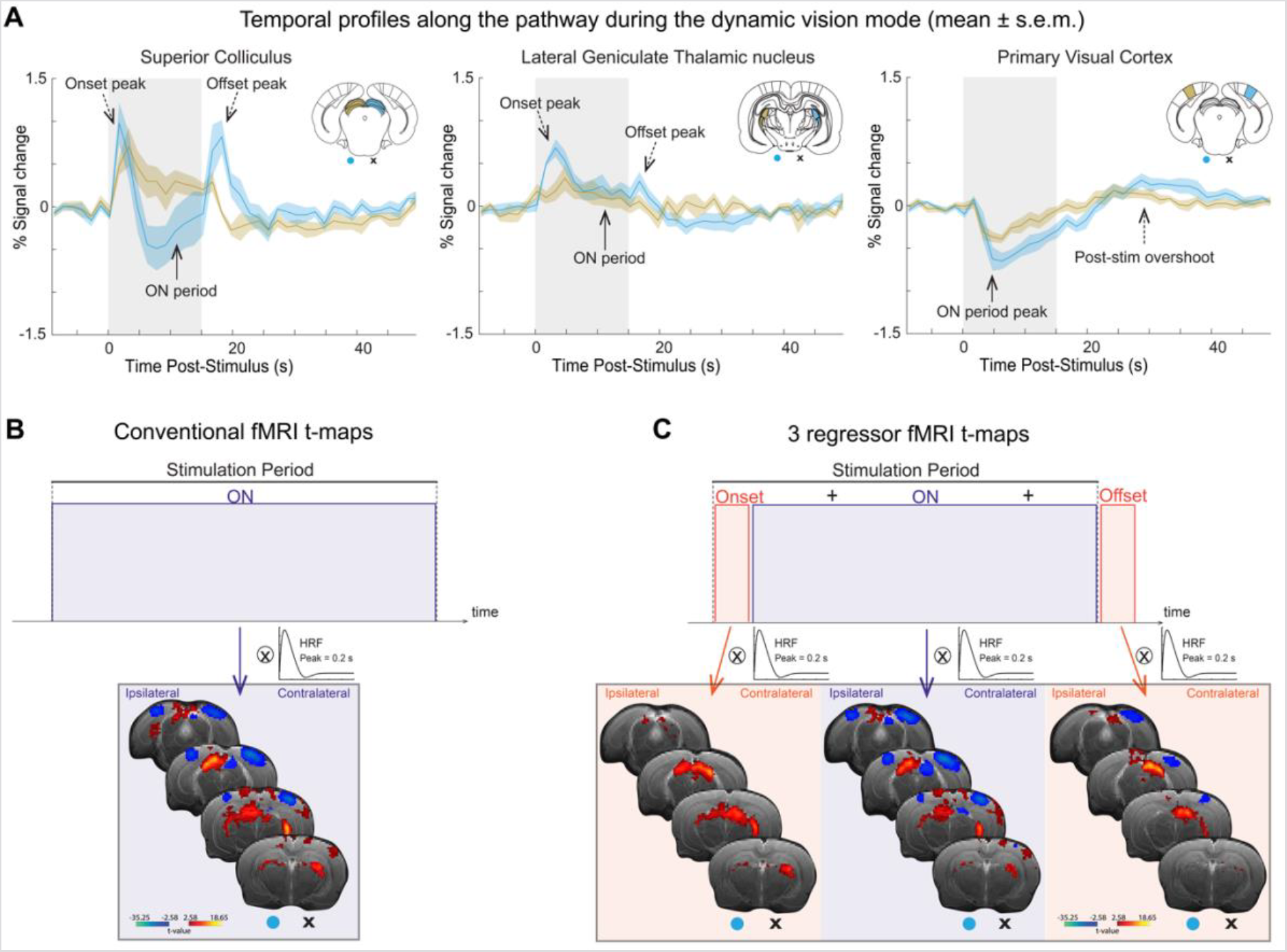
SC fMRI response for the healthy group during the dynamic vision mode. **(A)** Temporal profiles of the three main structures of the visual pathway: SC, LGN and V1. Responses for the contralateral and ipsilateral hemispheres are shown in blue and yellow, respectively. Contralateral multi-periodic responses can be observed in subcortical structures, particularly within the SC. For the SC and LGN, black arrows point to the 3 detected periods of the response: Onset peak (dashed arrow), ON period (full arrow) and offset peak (dashed arrow). For cortical responses ON period peak and post-stim overshoot are marked with filled and dashed black arrows, respectively; **(B)** FMRI t-maps assuming one regressor for the entire stimulation period. Negative cortical responses along with opposing tectal responses and positive contralateral LGN responses can be observed; **(C)** Onset and offset t-maps are highlighted in orange and the ON period t-map is highlighted in dark purple. The left side of the t-maps represents the ipsilateral hemisphere while the right side represents the contralateral hemisphere. T-maps were tested for a minimum significance level of 0.005 with a minimum cluster size of 10 voxels and corrected for multiple comparisons using a cluster false discovery rate test. SC - superior colliculus; LGN - lateral geniculate thalamic nucleus; V1 - primary visual cortex.

To understand the spatial distribution of the fMRI signals, a conventional GLM analysis tracking the whole block paradigm was first performed (**Figure 2B**). The corresponding t-maps revealed bilateral cortical negative fMRI activation centred in the V1, with a larger activation extent in the contralateral hemisphere. By contrast, at the subcortical level, opposing polarity SC responses were clearly observed: in the ipsilateral side, positive activation spanning the entire medial SC plane were noted, while the contralateral SC activation was clearly negative and appeared localised in more medial SC regions. In the LGN, positive activation covering the entire structure was observed, with higher t-values in the contralateral hemisphere.

The SC time profiles in **Figure 2A**, suggest that a multi-periodic analysis may be more relevant to decompose the activation in the onset, on, and offset epochs, respectively (**Figure 2C)**. Under this analysis, onset t-maps revealed that the onset peaks are localised only at the subcortical structures forming the parallel streams of the visual pathway - no activation could be detected for the onset signal in the cortical areas. The ON period t-maps strongly highlighted the opposite polarity of tectal responses: negative signals in cSC reaching a maximum t-value of about -35 and positive signals in iSC reaching maximum t-values of ∼18. These maps further confirm that contralateral responses cover more medial and dorsal regions of the cSC while the positive ipsilateral responses span the entire extent of the iSC. The ON period t-maps also demarcate the negative bilateral V1 responses during the dynamic vision mode. LGN ON signals are positive with a similar spatial distribution as LGN onset signals. Finally, offset t-maps show positive activation for subcortical structures, with the largest t-values in cSC. Particularly for the cSC, the offset signal spatial distribution covers the entire structure similarly to the onset t-maps. Offset signals in LGN appear to be more localised in the outer shell of this structure. Finally, negative signals appear for the V1 region which reflect the recovery of cortical signals to baseline.

### Diminished V1 feedback decreases cSC onset activation and ON suppression responses during the dynamic vision mode

To reduce corticotectal feedback, the V1 was bilaterally lesioned and experiments were repeated (**Figure 3**). The responses in both SC’s for this group are shown in **Figure 3A** (time profiles of the visual pathway structures can be seen in **Figure S2A**). In the cSC time profiles, the amplitude decreases were observed in the onset peak and ON periods. The cSC response (**Figure 3A**, solid blue line) was less negative compared to its control counterpart (**Figure 3A**, dashed blue line). The direction of the response is marked with a solid blue arrow in **Figure 3A**.

**Figure 3:**
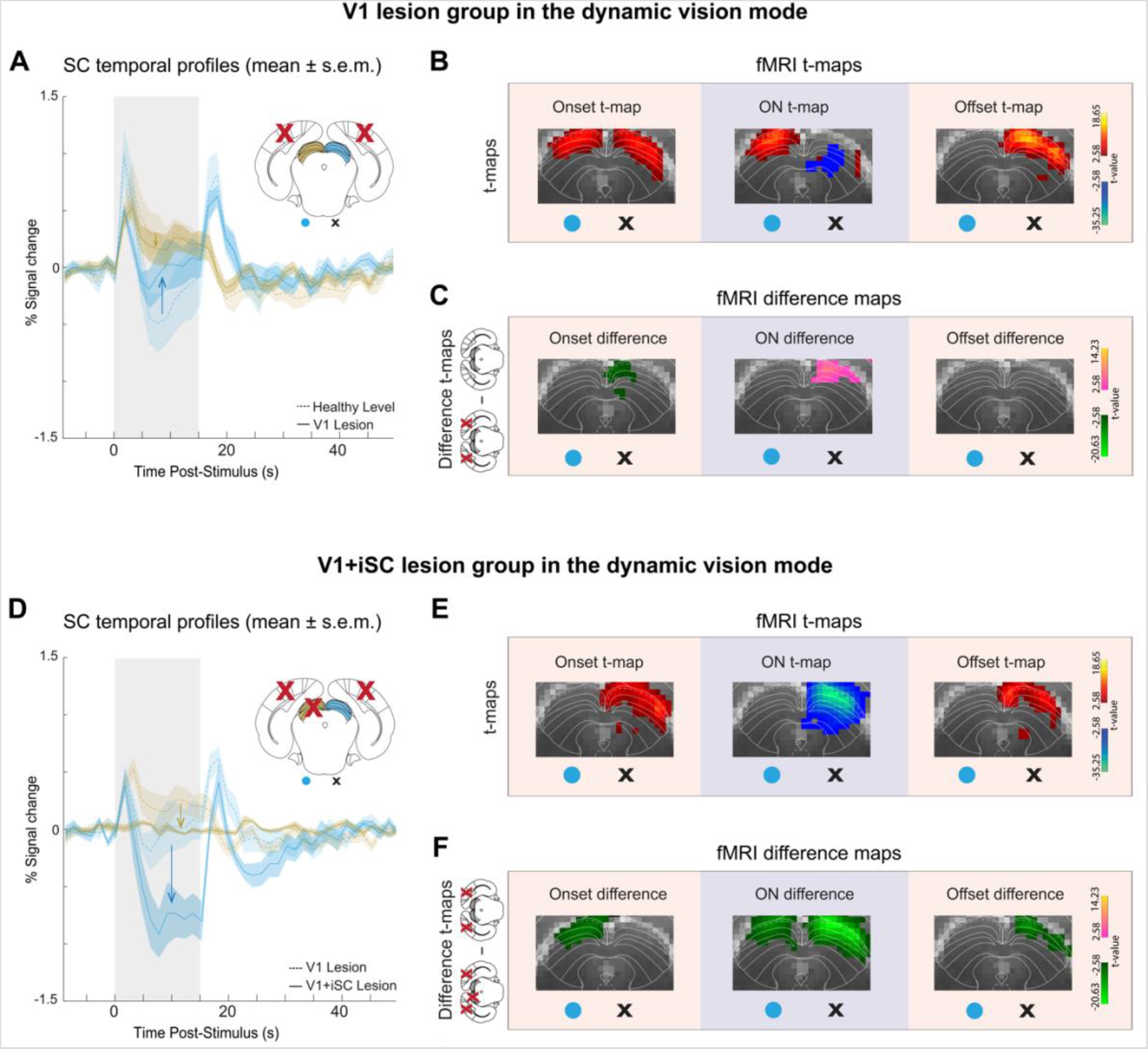
SC fMRI responses for the lesioned groups during the dynamic vision mode. **(A-C)** V1- lesioned group and **(D-F)** V1+iSC-lesioned group responses. **(A and D):** SC time profiles for the contralateral (blue) and ipsilateral (yellow) hemispheres. The healthy group and V1-lesioned responses are shown in dashed lines for clearer comparison in **A** and **D**, respectively; **(B and E)** Decomposition of the responses into the three detected periods and t-maps for each period. Onset and offset t-maps are highlighted in orange and the ON period t-map is highlighted in dark purple; **(C and F)** Difference maps for each response period. The left side of the maps represents the ipsilateral hemisphere while the right side represents the contralateral hemisphere. T-value maps were thresholded with p < 0.005 and a minimum cluster size of 10 voxels, and were cluster-FDR corrected at p < 0.005, both when analysing each group and the difference between groups. V1 - primary visual cortex; iSC - ipsilateral superior colliculus.

GLM maps calculated for the three onset, ON and offset periods, revealed interesting spatial distribution patterns (**Figure 3B** shows the main slice where most activation occurs for SC; other slices containing SC are shown in **Figure S2B**). Similarly to the healthy group (e.g. maps in **Figure 2C**), the V1-lesioned group exhibited strong positive onset activation in both SCs (**Figure 3B**). The opposing ON responses of the cSC and iSC, with weaker negative values in more medial cSC regions, and similarly positive values in the iSC areas, remained visible. Offset signals were confined to the cSC.

To obtain a more quantitative assessment of how cortical lesions affected activity in the tectum, difference t-maps were calculated for each signal period, representing the statistical differences (t-values) between the V1-lesioned group and the healthy animals (**Figure 3C**, other slices containing SC are shown in **Figure S2C**). In this fixed effects analysis, greener colours represent statistically negative differences between the groups, and warmer colours represent statistically positive differences. For both the onset and ON t-maps, a clear difference in t-values was observed in the more medial superficial cSC regions, with negative onset difference t-values (representing the decrease in the onset peak amplitude as seen in time profiles) and positive ON difference t-values (representing the less negative ON period amplitude after cortical lesioning). The offset signals showed no differences between the two groups. Similar experiments for the static vision mode are reported in **Figures S3 and S4**.

### Additionally lesioning iSC boosts cSC suppression in the dynamic vision mode

To dissect the role of intertectal connections in the push-pull interaction observed above and to avoid cortical loop effects, a third group was lesioned bilaterally in both the V1 (abolishing cortical feedback) and in the iSC (abolishing tectotectal feedback). **Figure 3D** shows ROI time profiles for this group (time profiles of all visual pathway structures can be seen in **Figure S2D**), represented by filled lines, revealed the expected flat ipsilateral tectal responses (**Figure 3D**, solid yellow line; the V1-lesioned response is shown for comparison in the dashed yellow line), confirming the effectiveness of the iSC lesion.

In the cSC, we find a starkly more negative response for the V1+iSC lesioned group (**Figure 3D**, solid blue line; the V1-lesioned response is in dashed blue for comparison). Interestingly, the onset peak is not markedly changed, while the offset peak is somewhat weaker than in the V1-lesioned group. The corresponding t-maps (**Figure 3E**, other slices containing SC are shown in **Figure S2E**) reveal that onset signals, similarly to offset signals, were circumscribed to the superficial cSC layers and spanned the entire medial plane of the structure. The ON t-maps evidenced strong negative values along the entire cSC and no positive iSC (lesioned) responses.

As above, we assess the differences more quantitatively via difference maps (**Figure 3F**, other slices containing SC are shown in **Figure S2F**). The decrease in the iSC responses was evidenced both in onset and ON difference maps. The onset maps reveal no differences in the onset cSC signals of both groups. The ON difference maps further revealed significantly more negative responses in cSC. Offset t-maps evidenced a small decrease in offset signals at the superficial cSC layers. Similar experiments for the static vision mode are reported in **Figures S3 and S4**.

## Discussion

The visual pathway switches between the static and dynamic vision modes, depending on the frequency of stimulus presentation, to encode for temporal stability and detect novel features in the surrounding environment. The rat SC, known for its involvement in RH^6,9–13^ and novelty detection^3,6–8^, has been recently implicated in the encoding of the transitions between these vision modes^15^. At frequencies lower than the FFF threshold, every stimulus elicits increased population level activity in the SC including elevated LFPs and multi-unit activity (MUA)^15,36,37^, and the system operates in the static vision mode. In particular, RH (characterised by a decrease of SC activity) progressively occurs in the SC as the inter-stimulus interval between light flashes decreases. In other words, stronger RH occurs as the stimulation frequency increases. As stimulation frequency increases beyond the FFF threshold, stimuli become fused and the neural activity in the SC is generally suppressed, entering the dynamic vision mode, with two exceptions: onset and offset signals clearly appear at the edges of the stimulation period, and were associated with the novelty of the initial transition from dark to bright in the very beginning of stimulation, as well as with the sharp transition from light back to darkness in the very end of the stimulation, each eliciting a novelty detection by SC^15,38^ (*vide infra*). This dual action, of encoding novelty by eliciting activity in SC and silencing SC beyond the FFF threshold, suggest potentially different SC modes of operation and potentially different interactions with other brain structures and even within the two superior colliculi. In the static vision mode, corticotectal projections are known to have a gain effect over the SC’s activity. Most studies suggested that these connections facilitate SC responses during the static vision mode^19,20,22–24,39^. Intertectal SC connections, associated with reduction of saccadic blur and visual-orienting behaviours through the “Sprague effect”^29^, have been mostly assumed to be inhibitory.

In the dynamic vision mode, less is known about the tectotectal and corticotectal interactions, motivating our current work. To investigate these interactions, we chose to harness fMRI due to its ability to noninvasively provide a whole-pathway overview of population level activity in the system (n.b. the excellent correlation between population-level activity - MUA - and fMRI signals observed recently under similar conditions^15,38^) along with brain lesions in bilateral cortical and unilateral SC areas that allow the abolishment of specific inputs into the SC. We further opted for monocular stimulation to isolate the role of each SC at a time.

Our findings suggest that, unlike its static vision mode counterpart, the dynamic vision mode evokes a marked push-pull interaction between the contralateral and ipsilateral SCs. As the system enters into dynamic vision mode, and the cSC activity is suppressed, the iSC engages in relatively strong activation. In other words, while the cSC “pushes” downward towards deactivation, the iSC “pulls” the signal upwards towards a less deactivated state. Indeed, when the iSC was lesioned, the iSC “pulling” was abolished, generating a dramatically stronger deactivation in the cSC (**Figures 3D-F**). In addition, the positive iSC response was quite independent of corticotectal feedback (as evident from the V1-lesioned group responses seen in **Figure 3A-C**) unlike its cSC counterpart (which is affected by corticotectal feedback), suggesting that the main push-pull interaction occurs at the tectotectal level, with little interference from other areas, potentially hinting at more complex tectotectal interactions than previously considered.

Based on these findings we can now propose a mechanism for the different interactions occurring during the dynamic vision mode (**Figure 4A**). SC activity is strongly suppressed when entering the dynamic vision mode and cortical feedback potentiates the suppression towards a maximal level (represented in green in **Figure 4A**): in binocular stimulation, the two SCs are suppressed to this level, likely signaling a bilateral agreement about the input. If, however the stimulation is monocular, the iSC activates, likely through tectotectal feedback from the cSC, exerting a “pull” effect on the cSC, leading to a less suppressed state, potentially either creating a diminished threshold for detecting novel stimuli in the eye that is seeing continuous light, or signalling that the two eyes are not in agreement.

**Figure 4:**
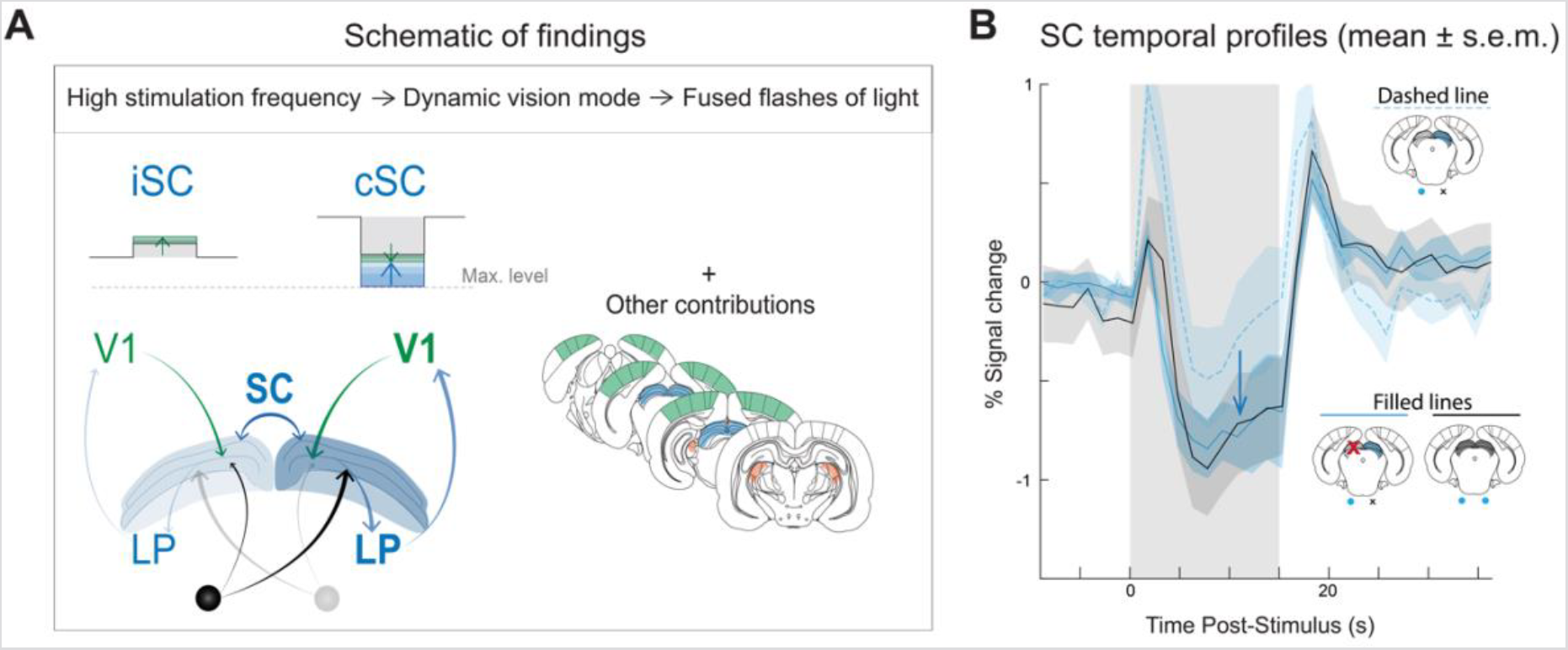
Proposed interactions during the dynamic vision mode. **(A)** The findings described in this study suggest a cortical amplification effect (represented in green) and a push-pull interaction between SC’s (represented in blue). As this study focused on the effects of corticotectal and tectotectal feedback and therefore, contributions from other visual pathway structures cannot be discarded. We propose that the high frequency stimulus (in grey), combined with the cortical potentiation effects (in green) induces maximal cSC suppression (and, consequently, maximal negative fMRI response) which is opposed by the positive iSC responses since only one eye is receiving such visual stimulus. Following this prediction, the cSC responses upon monocular stimulation would become similar to the negative SC responses upon binocular stimulation (where both SC’s would be maximally suppressed by the visual stimuli hitting both eyes combined with cortical potentiation) when the iSC is silenced; **(B)** When comparing the responses for the two stimulation regimes in healthy animals, the amplitude of the monocular responses (blue dashed line) showed less negative amplitude during the ON period than the binocular responses (black filled line). When the iSC was silenced (blue filled line) the monocular responses became identical to the binocular responses, supporting the counterbalancing role of the iSC on the negative cSC responses. cSC - contralateral superior colliculus; iSC - ipsilateral superior colliculus.

Given this proposed mechanism, we can make two simple predictions: (1) cSC responses upon monocular stimulation should present less negative responses compared to the (two) SC signals upon binocular stimulation (because the pull element is not present in the latter, both SC’s are both “cSC” and “iSC” in a sense); and (2) a lesion in iSC should prevent the tectotecal “pull” element exerted on the cSC in monocular stimulation, thereby allowing the cSC’s signal to reach the level it would potentially attain upon binocular stimulation signal in the dynamic vision mode. To test these predictions, we measured the SC signals under all these scenarios (**Figure 4B**). Indeed, our predictions are observed in these experiments. SC signals are more negative for binocular stimulation compared to monocular stimulation, and when we lesioned only iSC and performed monocular stimulation, the cSC signals became more negative, reaching the binocular stimulation levels. (**Figure 4B**, filled blue lines) when the iSC was silenced. We note in passing that our study focuses on the corticotectal and tectotectal feedback, and therefore contributions from other structures within the visual pathway, such as LP, LGN or higher order visual areas, cannot be ruled out. Nevertheless, our findings point to a major tectotectal push-pull effect that originates in the tectum.

The SC is part of the tectum which, additionally, encompasses two posterior inferior colliculi (ICs). The IC is a major midbrain auditory integration centre and as the SC and the IC are both tectal multimodal structures, they are indeed connected. For example, the IC coordinates with the SC in orienting the gaze toward or away from visual and auditory stimuli^5^ and an inferior-superior colliculus circuit providing key visual spatial attention signals has been reported^40^. Interestingly, a push-pull like mechanism^41–43^ - stronger contralateral excitation and ipsilateral inhibition - has been proposed to occur between the two ICs playing important roles in resolving sound intensity disparities in the context of sound localisation and lateralisation^41^. As such an effect in the IC has been associated with monaural and binaural sound integration, this SC push-pull interaction found here could be key in the encoding of lateralized fast-moving features within the field of view. Although this type of computations is usually associated with cortical areas^44^ in higher-order mammals, as already mentioned above, the rodent SC is more prominent for visual processing. Therefore, in lower-order animals, the involvement of SC in such visual localisation processing is plausible. However, future studies incorporating behavioural readouts are required to further elucidate on this matter.

The push-pull mechanism was clearly observed in the dynamic vision mode, but the static vision mode likely operates within a different interaction, as proposed in **Figure S5** based on the findings reported in **Figure S3**. Maximal positive responses in the cSC arise from a combination of the contralateral activity elicited by the visual stimulus (represented in grey in **Figure S5A**), cortical potentiation (represented in green in **Figure S5A**) and an intertectal boosting effect (represented in blue in **Figure S5A**), as both SCs present responses of the same polarity. Therefore, we hypothesise that this maximal activation regime would be reached both in healthy monocular cSC responses and in healthy binocular positive responses. Consequently, silencing iSC inputs would then result in less positive monocular cSC responses (**Figure S5A**). Indeed, the amplitude of healthy responses for both stimulation regimes (monocular and binocular) appeared similar, in-line with our hypothesis (**Figure S5B**), and a general decrease in positive amplitude of responses was seen in the monocular iSC-lesioned regime.

The distinct mechanisms proposed for the two vision modes (**Figure 4A and S5A**) suggest a higher degree of complexity at the commissural level between the two SCs than previously thought. Although intertectal SC connections have been mostly assumed to be inhibitory, reports of equal glutamatergic and GABAergic intertectal connections^30^ with similar topographic distribution in the cat support the idea that the tectotectal pathway may consist of two distinct functional components. Moreover, another study performed in cats, revealed specific spatial distributions of commissural excitation and inhibition, which were associated with saccadic eye movements: while excitation was recorded between the medial-medial or lateral-lateral parts of both SCs; commissural inhibition was observed between the medial SC on one side and the lateral SC on the opposite side^45^. Our work reveals that the suggested complexity of the tectotectal pathway also exists in the rat animal model for simple visual stimuli aiming at the entire animal field of view and where saccades are not expected to occur (therefore dissociating this effect from saccade occurrence). While further studies are needed, in particular integrating behavioural readouts, these changes in intertectal interactions during different vision modes (potentially activating different intertectal connections especially due to the distinct spatial distribution of positive and negative SC signals) may be important for instance in allowing adequate behavioural responses to different stimuli^30^ (for example, between slow and fast approaching objects). Interestingly, negative cSC responses were observed in more medial regions of the rat SC which have been associated with defensive mechanisms as these represent the upper visual field, from where most predators would attack. Due to the visual stimulus used in this study, which stimulated the entire field of view of animals, the interpretation of the distinct spatial distributions of the observed positive and negative fMRI tectal responses regarding their behavioural relevance is not straightforward; however, it is nonetheless curious to note that contralateral negative responses during the dynamic vision mode appear more colocalized in regions associated with defensive type of behaviours.

The above discussion has focused mostly on the observed ON period modulations of SC responses; however, another important finding of our work is that fMRI signals contain “fine structure” that allows for detection of response dynamics, including onset and offset signals (associated with novelty detection). It is interesting to notice how cortical gain did not play a significant role on offset responses (evidenced by the difference maps in **Figure 3C**), while tectotectal input mostly affected these signals (evidenced by the difference maps in **Figure 3F**). This suggests that the two signal peaks could be underpinned by different sources, and these could be studied in the future e.g. with electrophysiology. A limitation of the fMRI “fine structure” is that in some areas that do not exhibit such features, multiple effects could be conflated upon a GLM, and care is needed for instance not to mistake a post-stimulus (over)undershoot with an offset signal. Nevertheless, this kind of decomposition may be beneficial in certain contexts, for instance, when additional electrophysiology has been performed and the signals are confirmed as onset/offset as in our recent work^15^.

Finally, several methodological aspects and limitations merit further discussion. In the current study, we resorted to brain lesions targeting key structures of the visual pathway to selectively reduce tectal feedback connections. One disadvantage of using such a method for silencing a specific region is the variability of the lesions performed in different animals. While successfully silencing the desired region, the specific coverage of each lesion or the inflammation still present after a certain time interval, may slightly vary from animal to animal. Furthermore, lesioning a brain region can lead to neural plasticity events that may already take place during the period between the lesion and data acquisition. In the current study lesioned animals were allowed to recover for 7-9 days before being imaged. This time interval was chosen to allow for the animal’s recovery, for the lesions to produce their effects, and to minimise inflammation (which on its own could be a confounding effect). Although at least a partial recovery of SC’s normal activity is expected according to previous studies^46^, this recovery process probably also involves some degree of plastic changes in other areas and connections, in an attempt to compensate for the initial lesion. Larger plasticity events however, are thought to take place across longer time frames^46^. Importantly, our results are qualitatively comparable to Goodale’s work^18^, which included collicular and cortical lesions in the rat during bipolar electrode recordings and where activity was recorded immediately before and after the lesion. This similarity points at a relatively small contribution of plasticity in our measurements. Future studies harnessing e.g. optogenetics^47–50^ or pharmacological^51,52^ studies for silencing specific areas may provide additional insights in future studies.

Moreover, the use of anaesthesia when studying functional processes may affect the interplay between bottom-up and top-down pathways differently depending on the anaesthetic used. Particularly for the SC, it has been shown that under the effect of urethane^18,53^ or isoflurane^21,54^, the corticotectal connections, top-down inputs to the SC, are more affected than the bottom-up retinal inputs or the intertectal interactions, and can sometimes even be completely masked^19^. In the present study animals were lightly sedated with medetomidine, an α2 agonist^55^, whose effects have been shown to be small when comparing baseline neural activity to awake^56^. A good agreement between SC responses under medetomidine sedation and behavioural measurements have been observed^15^ along with similarities to studies in awake mice^57^. Furthermore, studies in rats, with a similar medetomidine protocol as the one used in this work, have reported functional connectivity which was partially attributed to wakefulness^58^. Although we cannot rule out possible changes in the visual pathway during medetomidine sedation, taking these favourable comparisons into account, we believe that the use of medetomidine anaesthesia did not strongly affect our results.

## Conclusion

We report a push-pull interaction between the two SCs in the dynamic vision mode where individual flashes are perceived as a continuous light stimulus. While cSC responses were suppressed by the visual stimulus, positive responses in the iSC pulled cSC into a less deactivated state, and these interactions were observed independently of cortical lesions. Solely silencing the iSC responses led to stronger negative cSC responses identical to the healthy binocular responses observed during the dynamic vision mode. The proposed interactions occurring between the two SCs were different for low and high stimulation frequencies suggesting a complex SC intertectal interplay that is modulated by stimulation frequency. Finally, the distinct spatial segregation of positive and negative SC responses suggests different tectotectal mechanisms in the static vs. the dynamic vision mode, each probably associated with activation of different commissural connections. Our findings suggest a higher complexity of the tectotectal pathway than a simple reciprocal inhibitory effect.

