## Supplementary Information for "Evidence for a push-pull interaction between superior colliculi in monocular dynamic vision mode"

One Sentence Summary: Opposing signals between superior colliculi in the dynamic vision mode suggest an active push-pull interaction within the tectotectal commissural pathway.

### The PDF file includes:

- Supplementary Results and Figures:
    - **Figure S1:** Raw data and Averaged time profiles
    - Dynamic vision mode:
        - **Figure S2:** FMRI time profiles for other visual pathway structures and maps of other brain slices that contain the SC during the dynamic vision mode.
    - Static Vision mode:
        - Discussion on Cortical and ipsilateral tectal potentiation of contralateral positive responses exist during the static vision mode
        - **Figure S3:** Visual pathway responses during the static vision mode
        - **Figure S4:** FMRI time profiles during the static vision mode
        - **Figure S5:** Proposed interactions during the static vision mode

### Supplementary Figures:

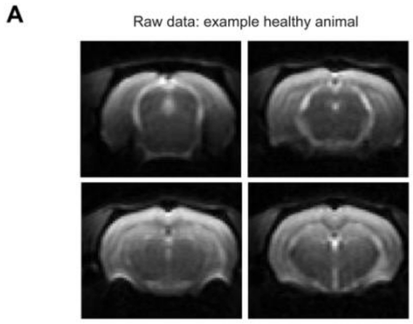


**B**

tSNR table

| Groups \ ROIs | Contralateral SC (mean ± s.e.m) | Contralateral LGN (mean ± s.e.m) |
|---|---|---|
| Healthy Group | 45.8 ± 3.9 | 35.0 ± 2.6 |
| V1-lesioned Group | 39.4 ± 2.6 | 27.8 ± 2.1 |
| V1+iSC-lesioned Group | 40.6 ± 0.6 | 31.8 ± 2.2 |
| iSC-lesioned Group | 36.0 ± 3.1 | 30.1 ± 2.8 |

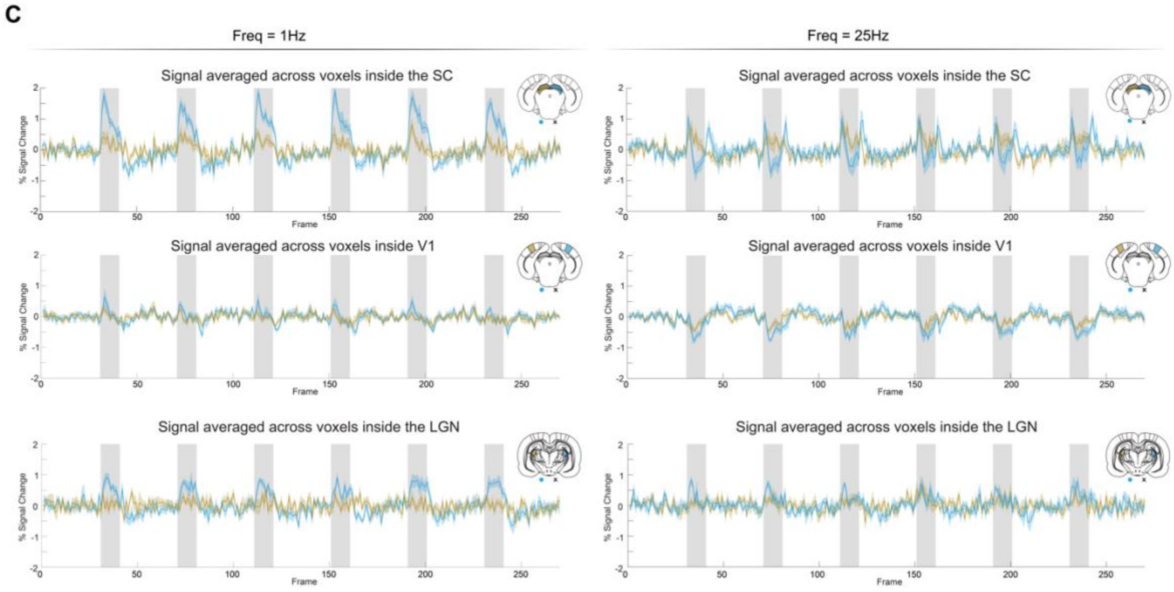


**Figure S1: Raw data and averaged time profiles. (A)** Raw functional brain images from one example healthy animal revealing the good data quality; **(B)** Table showing the tSNR values calculated in the cSC and cLGN for the different groups in the study. The similar values obtained between the groups revealed that the SC response modulation observed did not originate from differences in tSNR; **(C)** Averaged fMRI runs for the main structures of the visual pathway in the healthy group. Profiles obtained for a low stimulation frequency of 1 Hz, inducing the static vision, are shown on the left side while profiles obtained for a high stimulation frequency of 25 Hz, inducing the dynamic vision mode, are shown on the right side. Blue and yellow profiles represent the contralateral and ipsilateral hemispheres, respectively. Clear activation/suppression of fMRI responses can be seen along the six blocks of stimulation with no evidence of habituation. The different periods - onset, ON and offset - of the cSC responses are also observed for every block of stimulation. tSNR - temporal signal-to-noise ratio; cSC - contralateral superior colliculus.

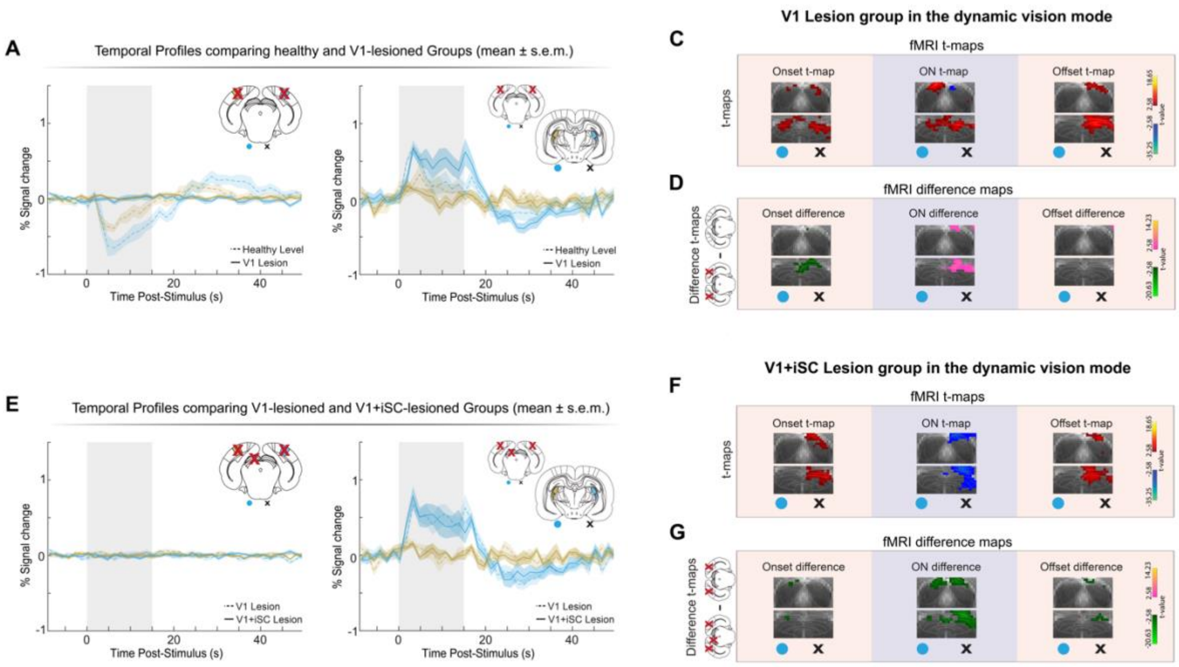


**Figure S2: FMRI time profiles for other visual pathway structures and maps of other brain slices that contain the SC during the dynamic vision mode. (A)** Time profiles for the V1-lesioned group (with healthy responses in dashed lines for easier comparison). Cortical profiles are flat since lesions were performed in both V1s and contralateral thalamic responses appear positively modulated upon cortical lesions; **(C-D)** Maps shown in **Figure 3B-C** for other acquired brain slices encompassing the SC. The maps reveal a homogeneous response profile along the A-P extension of the structure; **(E)** Time profiles for the V1+iSC-lesioned group (with V1-lesioned responses in dashed lines for easier comparison). Cortical profiles are flat since lesions were performed in both V1s and contralateral thalamic responses do not change with further iSC lesions; **(F-G)** Maps shown in **Figure 3E-F** for other acquired brain slices encompassing the SC. The maps reveal a homogeneous response profile along the A-P extension of the structure. T-value maps were thresholded with $p < 0.005$ and a minimum cluster size of 10 voxels, and were cluster-FDR corrected at $p < 0.005$, both when analysing each group and the difference between groups. V1 - primary visual cortex; iSC - ipsilateral superior colliculus.

### Cortical and ipsilateral tectal potentiation of contralateral positive responses exist during the static vision mode

**Figure S3** shows the results of cortical and ipsilateral tectal lesion during the static vision mode. In **Figure S3A** time profiles reveal general positive responses (stronger in the contralateral hemisphere). These time profiles reveal how the signals in the low frequency regime are not multi-periodic and therefore do not require a three-period decomposition, as in the high frequency regime. Conventional fMRI t-maps with one regressor representing the entire simulation period convolved with the chosen HRF, reveal cortical positive responses in secondary visual regions and more anterior V1 regions. The posterior V1 region (where negative responses are observed during the dynamic vision mode) did not show clear activation. Furthermore, positive responses along the SC and LGN appear homogeneous and span the entire visual area of these structures. Ipsilateral positive SC and LGN responses are also observed but with smaller t-values.

Bilateral lesions in V1 (**Figure S3B**) led to a reduction of SC signal amplitude for both hemispheres. Difference maps comparing responses in the healthy and V1-lesioned group revealed an overall signal reduction at the level of the superficial layers of both SCs. Further lesioning the iSC (**Figure S3C**), resulted in additional reduction of tectal responses. FMRI t-maps reveal contralateral subcortical activation and the difference map between the V1 and V1+iSC lesioned conditions revealed decreases in tectal responses concentrated in the superficial layers of the SC. T-value maps were thresholded with $p < 0.005$ and a minimum cluster size of 10 voxels, and were cluster-FDR corrected at $p < 0.005$, both when analysing each group and the difference between groups.

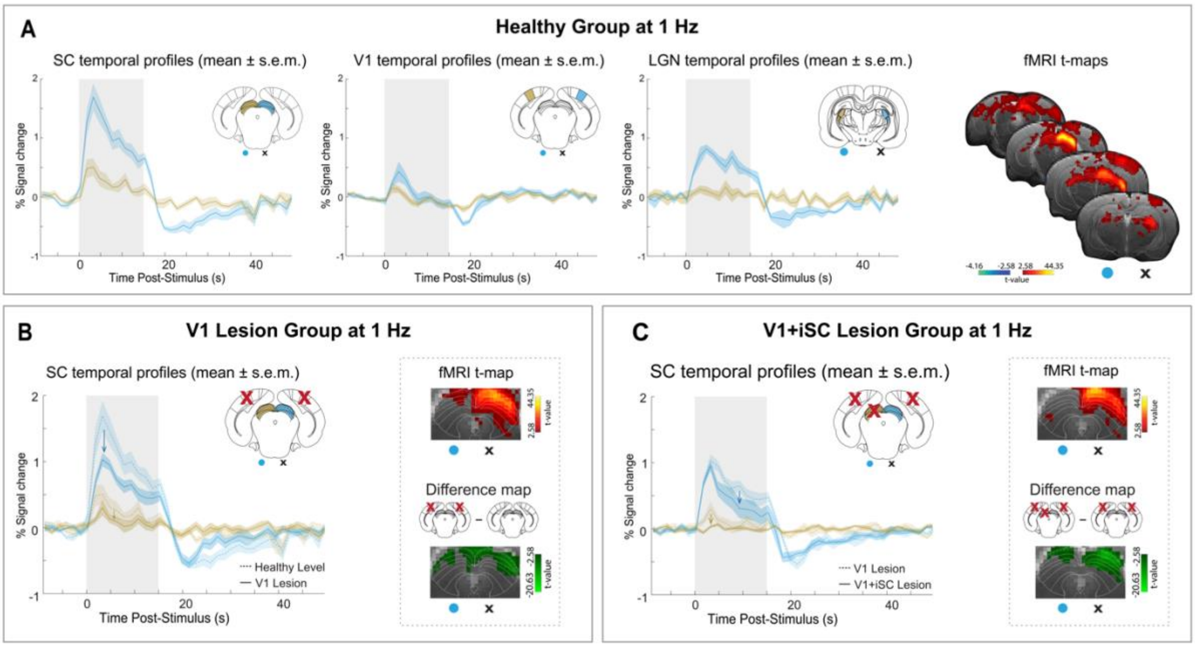


**Figure S3: Visual pathway responses during the static vision mode. (A)** Healthy group time profiles. Contralateral responses in the three main visual pathway structures reveal increased percent signal change compared to ipsilateral responses, as expected from monocular stimulation. T-maps confirm the positive responses along the visual pathway with larger t-values in the contralateral hemisphere; **(B)** V1-lesioned group responses. SC responses show decreased signal amplitude due to the cortical lesions in both hemispheres. Healthy responses are plotted in dashed lines for easier comparison between groups. T-maps reveal positive cSC and weaker medial iSC responses. The difference map between the V1-lesioned and healthy groups evidences the response reduction in both SCs superficial layers. **(C)** V1+iSC-lesioned group responses. SC time profiles evidence contralateral response decrease after iSC input was silenced due to the lesion. iSC profiles appear flat as expected due to the brain lesion. V1-lesioned profiles are shown in dashed lines for better comparison between groups. T-maps show similar results compared to the V1-lesioned group in the contralateral hemisphere. The difference map between V1+iSC-lesioned and V1-lesioned groups evidences the decrease in tectal responses. T-value maps were thresholded with $p < 0.005$ and a minimum cluster size of 10 voxels, and were cluster-FDR corrected at $p < 0.005$, both when analysing each group and the difference between groups. V1 - primary visual cortex; iSC - ipsilateral superior colliculus.

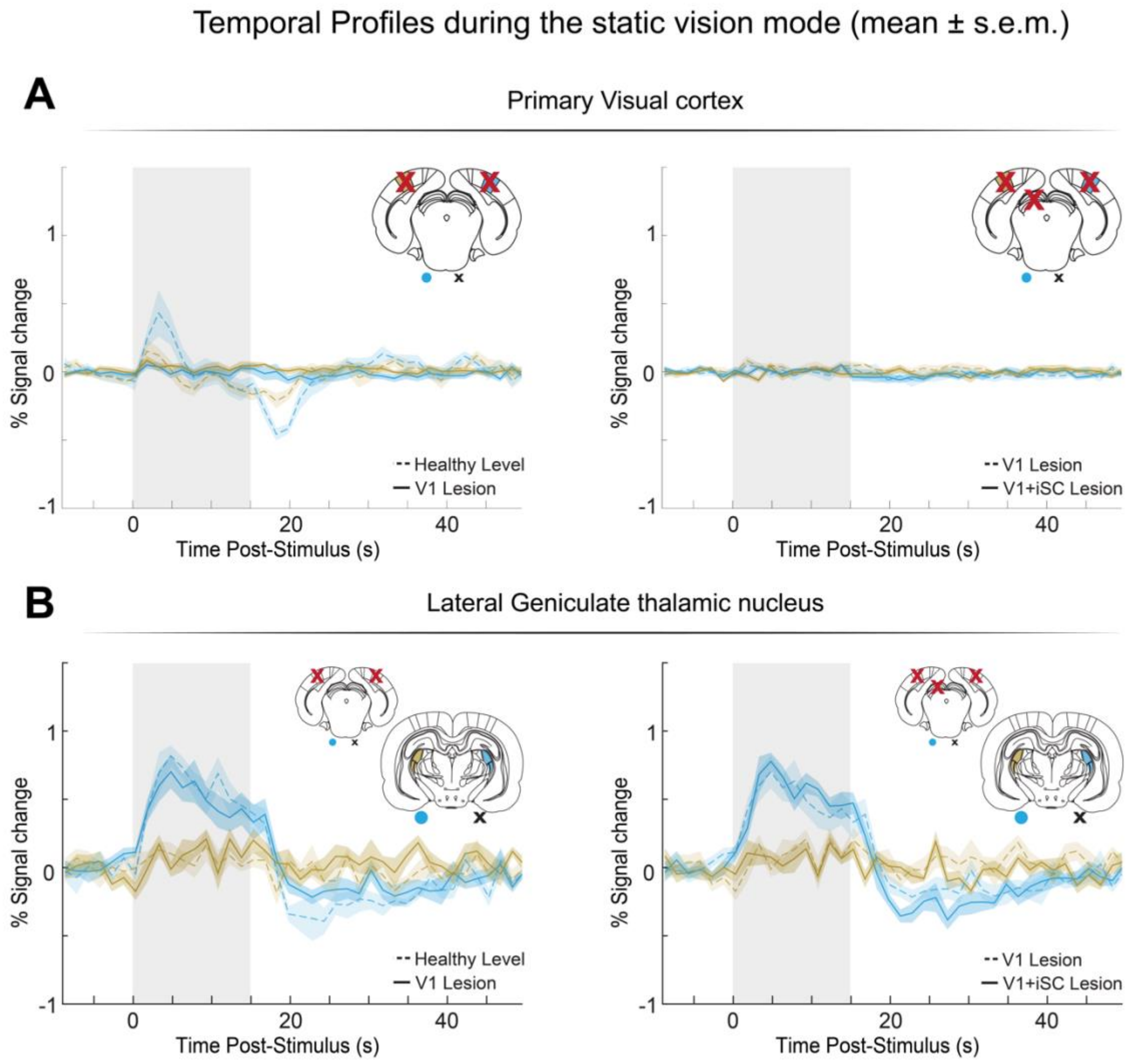


**Figure S4: FMRI time profiles during the static vision mode. (A)** Primary visual cortex and **(B)** Lateral geniculate thalamic nucleus. Left side plots reveal responses for the V1-lesioned group (with healthy responses in dashed lines for easier comparison) and right-side plots reveal responses for the V1+iSC-lesioned group (with V1-lesioned group responses in dashed lines for easier comparison). Cortical profiles are flat for both lesioned groups since lesions were performed in both V1s. Contralateral thalamic responses appear positive and remain similar between the different animal groups. No clear ipsilateral responses are observed in the LGN. V1- primary visual cortex; iSC - ipsilateral superior colliculus, LGN - lateral geniculate nucleus of the thalamus

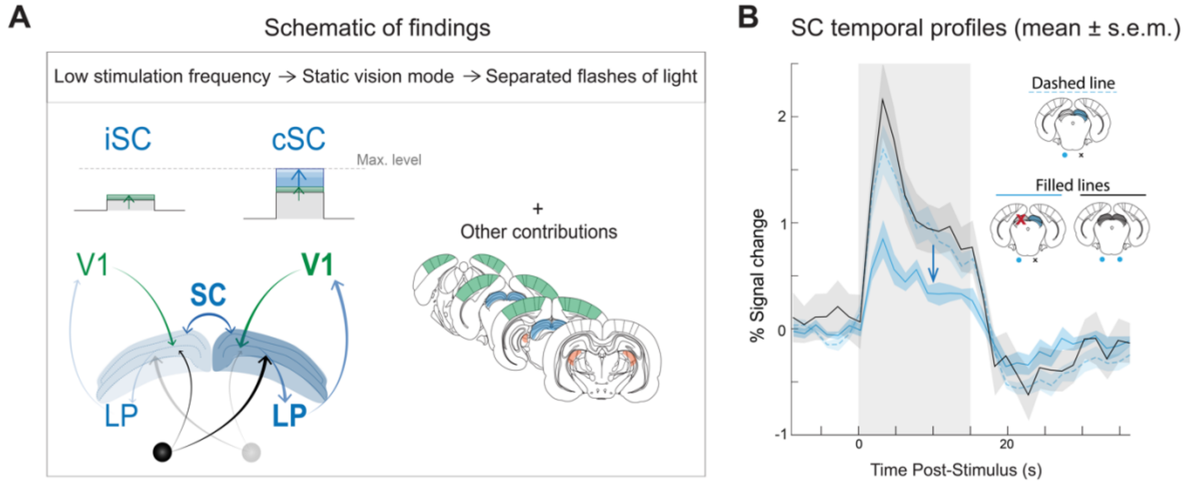


**Figure S5: Proposed interactions during the static vision mode. (A)** A cortical amplification effect (represented in green) and a boosting effect of the iSC on cSC (represented in blue) are suggested. This study focused on the effects of corticotectal and tectotectal feedback and therefore, contributions from other visual pathway structures cannot be discarded. We propose that the low frequency stimulus induces cSC activation (and, consequently, positive fMRI response) which is boosted by the iSC and cortical amplification reaching maximal positive values. Following this prediction, the cSC responses upon monocular stimulation would become similar to the positive SC responses upon binocular stimulation (where both SC's would be activated up to their maximal level by the visual stimuli hitting both eyes alongside the boosting cortical and intertectal effects) in healthy conditions; **(B)** When comparing the responses for the two stimulation regimes in healthy animals, the amplitude of the monocular responses (blue dashed line) show identical positive amplitude during the ON phase as the binocular responses (black filled line). When the iSC is silenced (blue filled line) the responses become reduced, supporting the prediction of intertectal boosting effects when both SCs present positive fMRI responses.
